# Multiplexed End-point Microfluidic Chemotaxis Assay using Centrifugal Alignment

**DOI:** 10.1101/2020.07.09.195644

**Authors:** Sampath Satti, Pan Deng, Kerryn Matthews, Simon P. Duffy, Hongshen Ma

## Abstract

A fundamental challenge to multiplexing microfluidic chemotaxis assays at scale is the requirement for time-lapse imaging to continuously track migrating cells. Drug testing and drug screening applications require the ability to perform hundreds of experiments in parallel, which is not feasible for assays that require continuous imaging. To address this limitation, end-point chemotaxis assays have been developed using fluid flow to align cells in traps or sieves prior to cell migration. However, these methods require precisely controlled fluid flow to transport cells to the correct location without undesirable mechanical stress, which introduce significant set up time and design complexity. Here, we describe a microfluidic device that eliminates the need for precise flow control by using centrifugation to align cells at a common starting point. A chemoattractant gradient is then formed using passive diffusion prior to chemotaxis in an incubated environment. This approach provides a simple and scalable approach to multiplexed chemotaxis assays. Centrifugal alignment is also insensitive to cell geometry, enabling this approach to be compatible with primary cell samples that are often heterogeneous. We demonstrate the capability of this approach by assessing chemotaxis of primary neutrophils in response to an fMLP (N-formyl-met-leu-phe) gradient. Our results show that cell alignment by centrifugation offers a potential avenue to develop scalable end-point multiplexed microfluidic chemotaxis assays.

## Introduction

Cells have a sophisticated ability to sense gradients of chemoattractants and then respond by directed migration along these gradients via chemotaxis. Chemotaxis underpins a diverse range of biological processes, including infection^1^, wound healing^2^, inflammation^3^, embryogenesis^4^, and cancer metastasis^5^. Consequently, there has been a longstanding interest to develop assays for chemotaxis. Conventional chemotaxis assays are often qualitative and low-throughput. Overcoming these limitations is key to integrating chemotaxis into platforms for diagnosis^6^, drug testing^7,8^ and drug discovery^9^. The traditional approach for evaluating chemotaxis uses the Boyden assay, also known as the Transwell assay, which generate chemical gradients across a porous membrane^10^. The cells that migrate across the membrane are then enumerated to produce a measure of chemotactic capability. While this approach is simple to perform, it often produces inconsistent results because the chemical gradient experienced by each cell across the membrane interface can vary with time, location, and cell density^9^. Transwell assays also require a large number of cells (10^5^ - 10^6^) and large volumes of chemoattractant solutions, limiting the number and types of experiments that could be performed, especially on primary cell samples obtained from patients^11^.

Microfluidic approaches have been used to develop improved chemotaxis assays that provide more reliable chemical gradients, as well as that reduce the requirements for cells and reagents^12–20^. These devices typically involve generating chemoattractant gradients through passive diffusion or through active perfusion. Cells are then introduced into this gradient in order to observe their migration via time-lapse microscopy. This approach has a limited capacity to perform multiple chemotaxis assays in parallel because of the need to continuously track cell migration in each assay using microscopy. While multiplexing these experiments could be achieved using automated microscopy, this approach is not scalable for high throughput applications such as drug testing and drug screening^21^.

To address the need for scalable multiplexed chemotaxis assays, several approaches have been developed to align the cell sample at a common starting point before starting chemotaxis experiments^22–25^. These approaches include trap-based alignment and sieve-based alignment. Trap-based alignment techniques direct cells into single-cell sized traps insider a larger microchannel using flow. Sieve-based alignment flow cells to the entrance of microchannels with a thickness smaller than the cell diameter (typically 5-10 µm). Since fluids can flow into these microchannels, but cells cannot, the cells are aligned at the entrance of these channels. A key limitation in the scalability of both trap- and sieve-based cell alignment is the requirement for precisely controlled fluid flow in each device to seed cells in the right locations. The flow rate must be matched with the cell type and size in order to align the cells without squeezing past the trap or sieve, or apply shear stress that may affect chemotactic behavior^26^. Consequently, this approach requires lengthy set up time for cell alignment and introduces design constraints that limit the scalability of the assay.

Here, we developed a scalable end-point chemotaxis assay that uses centrifugal force to align cell samples to a common starting point in order to provide a multiplexed end-point chemotaxis assay without the need for continuous microscopy. Individual microfluidic devices are arrayed in a rotational symmetric manner on a glass slide substrate. Unlike trap- or sieve-based alignment, this approach obviates the need for carefully controlled fluid flow by aligning cells via centrifugation against a barrier feature. We validated this approach by assessing the migration of primary neutrophils along an fMLP gradient in order to demonstrate its potential as a simple, scalable, and multiplexed chemotaxis assay requiring minimal equipment.

## Results

### Microfluidic Device Design and Assay Preparation

The microfluidic device for end-point chemotaxis assays comprise of two reservoirs connected by a thin microchannel that limits diffusion between the reservoirs (**Figure 1A & 1C**). The microfluidic channel contains a barrier feature that enables centrifugal alignment of the cell sample at a common starting point before starting each chemotaxis assay (**Figure 1B**). The barrier feature includes a small degree of concavity designed to trap cells against movement created by tangential forces (perpendicular to channel) resulting from angular acceleration caused by starting and stopping the spinning process. The microfluidic channel and alignment barrier are formed using PDMS bonded to a glass substrate, which can be functionalized with extracellular matrix (ECM) to enhance chemotaxis. To multiplex chemotaxis experiments, individual devices are arrayed in a rotationally symmetric fashion on a single glass substrate allowing for multiple experiments to be performed simultaneously (**Figure 1D**). As a demonstration of this approach, we developed a prototype multiplexing 12 separate devices on a single 50 x 75 mm glass slide substrate.

**Figure 1.**
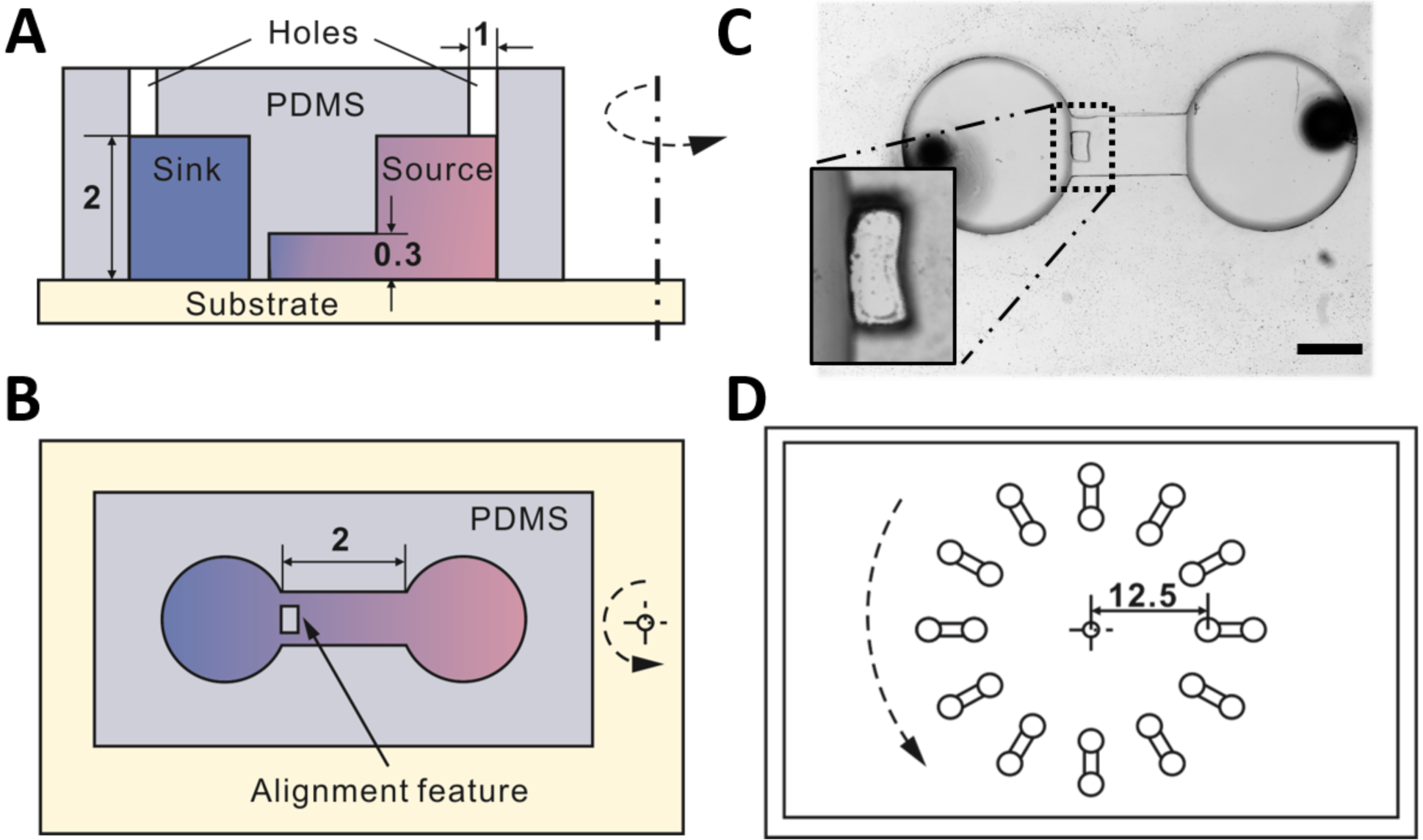
Design of the multiplexed end-point chemotaxis assay. **(A-B)** Cross-section and top-view schematics of the microfluidic device design for an individual chemotaxis assay, including the alignment barrier and critical dimensions (in mm). Chemoattractant gradients are generated by passive diffusion between the source and sink reservoirs. **(C)** Micrograph of an individual chemotaxis assay. Scale bar = 1000 µm. **Inset:** Concave barrier for cell alignment. **(D)** Design of the 12-plex end-point chemotaxis assay where cell alignment is achieved through centrifugal alignment. Radial position of the alignment feature in mm.

The multiplexed end-point chemotaxis assay begins by functionalizing the glass surfaces in each microfluidic device using fibronectin, an ECM protein that enhances neutrophil migration *in vitro*^27^. The devices are then blocked using bovine serum albumin (BSA) to prevent non-specific adhesion. After functionalizing the surfaces, the cell samples are introduced into each device by standard pipette. Next, the microfluidic device is spun in order to align cells in the device against the barrier feature by centrifugation. Cells not aligned against the barrier feature are transported to the reservoir past the barrier feature and generally excluded from the assay. In order to prevent fluid leakage during the alignment process, the outlets were sealed using acrylic tape. After pipetting the cell sample, the chemoattractant was pipetted into the source reservoir. Chemoattractant molecules were diffused from the source reservoir into the sink reservoir through the gradient channel to establish a gradient (**Figure 1B**). The device was then placed inside an incubator to allow chemotaxis to take place at stable temperature and gas conditions. Finally, after an appropriate amount of time, the device is imaged to determine the resulting cell positions.

### Gradient Formation

To study gradient formation in our microfluidic device, as well as the impact of the barrier feature, we first developed finite element models of the gradient with and without a barrier feature. Our model showed that the barrier feature caused the gradient profile to flatten near the barrier, which likely results from reduced fluid exchange in this area, but produced the expected experimental profile away from the barrier (**Figure 2A**). We then performed diffusion experiments using 10 kDa FITC-Dextran to visualize the gradient profile (**Video S1**). Our results show gradient profiles of both devices can be established after 30 minutes and maintains a stable value from 60 to 180 minutes (**Figure 2B-D**). Similar to our simulation results, our experimental results show that there is a flattening of the gradient profiles near the barrier feature compared to the microchannel without the barrier feature. Specifically, the gradient strength measured at the barrier feature (x = 0 in the Figure 2C inset) is smaller by a ratio of 0.62 compared to a microchannel without the barrier feature (**Figure 2D**). The correlation coefficient between gradient strengths in these two scenarios is 0.96, suggesting that the difference is a simple scale factor error.

**Figure 2.**
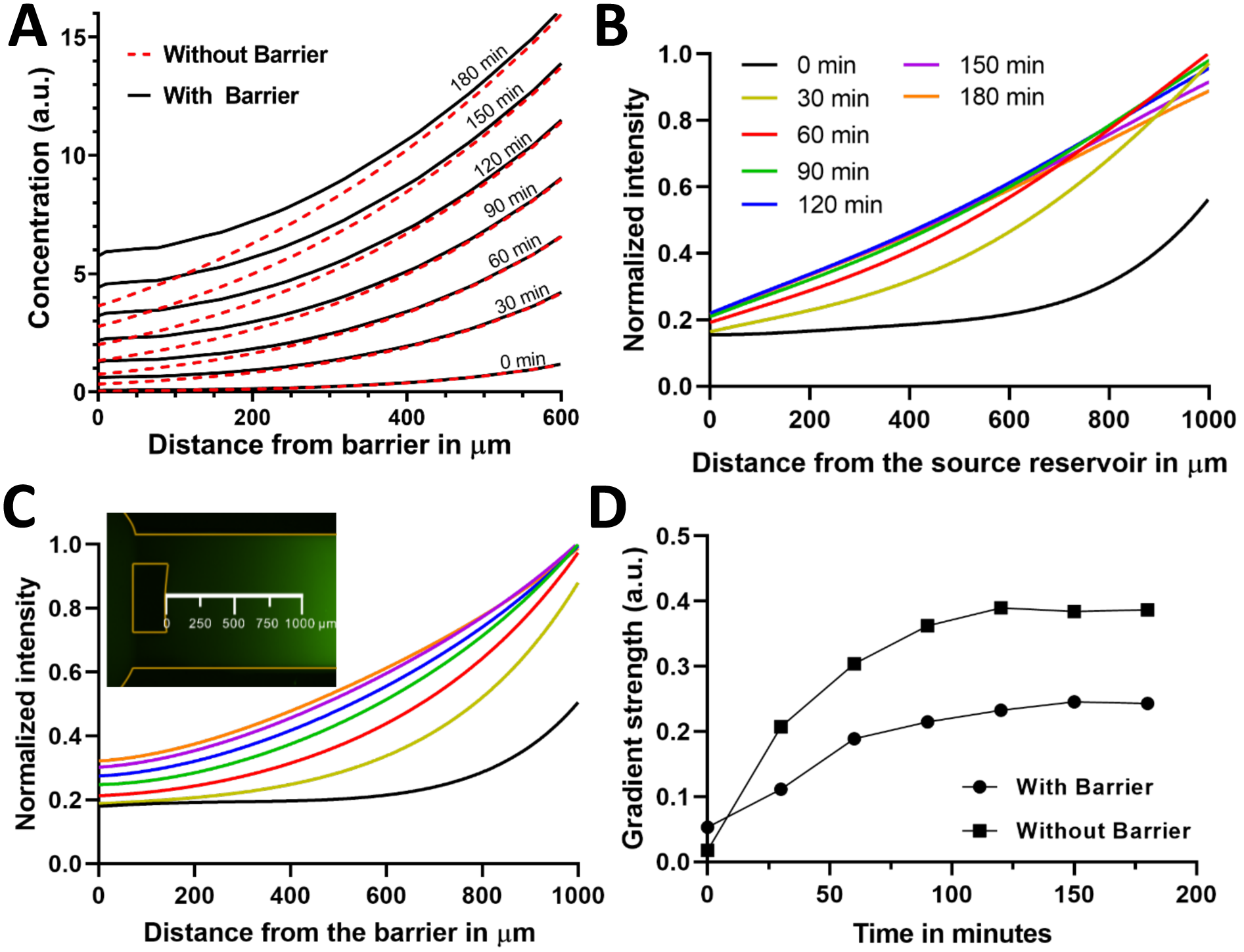
Simulation and experimental study of gradient generation using passive diffusion. (**A)** Finite element simulation of the gradient profile with and without the alignment barrier from 0 to 180 minutes. (**B**-**C**) Experimental study of gradient profiles without the barrier feature (B) and with the barrier feature (C). The gradient profile was visualized using fluorescent FITC-Dextran (inset in C). (**D)** The gradient strength measured at x = 0 µm (defined in inset in C) as a function of time for microfluidic devices with and without the barrier feature (correlation coefficient >0.96).

To ensure the gradient profile formed using FITC-Dextran is representative of the gradient profile formed using fMLP, we further tested gradient formation using Rhodamine B, which has a similar molecular weight as fMLP (Rhodamine B MW = 442.55 Da, fMLP MW = 437.55 Da). The Rhodamine B gradient is similar in profile to the FITC-Dextra gradient with a scale factor of 1.59 and a correlation coefficient of 0.99 (**Figure S1**). This scale factor error can be accounted for by modulating the chemoattractant concentration at the source reservoir.

### Cell Alignment Through Centrifugation

To align cells against the barrier feature, the microfluidic device is rotated using a spinner to produce a centrifugal force (*F*_*c*_) towards the center of rotation:

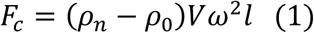

where *ρ*_*n*_ is the density of neutrophils^28^ and *ρ*_0_ is the density of water, *V* is the volume of a neutrophil, *ω* is the angular velocity, *l* is the distance from the axis of rotation. For small particles moving in a liquid at low Reynolds number, they need to overcome the Stokes drag force (*F*_*d*_):

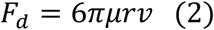

where *μ* is the dynamic viscosity of the fluid, *r* is the radius of small particles or cells, *v* is the speed of small particles or cells. Since the bottom surface of the microchannel is treated with fibronectin, *F*_*c*_ need be sufficiently large to first allow cells to overcome fibronectin adhesion before transporting cells against viscous forces towards barriers. Once cells begin to move, this adhesion force becomes negligible. Additionally, since the length the cells need to move across the microchannel is much smaller than *l*, the value of *F*_*c*_ could be considered constant during transport by centrifugation. When *F*_*c*_ is balanced by *F*_*d*_, the centrifugal transport velocity (*V*_*c*_) becomes:

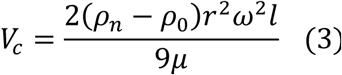

If cells could migrate consistently at *V*_*c*_ and cells were evenly distributed in channels at the beginning, the number of collected cells at barriers (*N*) could be calculated by:

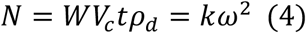

Where *W* is the width of migration channel, *t* is the centrifugal spinning time, *ρ*_*d*_ is the cell distribution density, *k* is a constant.

To experimentally confirm our model, we loaded the microfluidic device with human neutrophils (1.4×10^6^/ml) and centrifuged the device at speeds from 0 to 2000 RPM for one minute and then counted the number of collected cells (n = 5, **Figure 3A-D**). We found the number of collected cells to monotonically increase from 0 to 1500 RPM. At 2000 RPM, however, the number of collected cells decreased due to excessive tangential forces arising from spinner acceleration and deceleration. This issue could be resolve in future versions using a larger concavity. We then measured the value of *k* by fitting Equation (4) to data points from 0 to 1500 RPM and found a high-quality fit (*R*^2^> 0.98, **Figure 3E**). Based on these results, we selected 1500 RPM for 1 min as the condition for cell alignment throughout our study.

**Figure 3.**
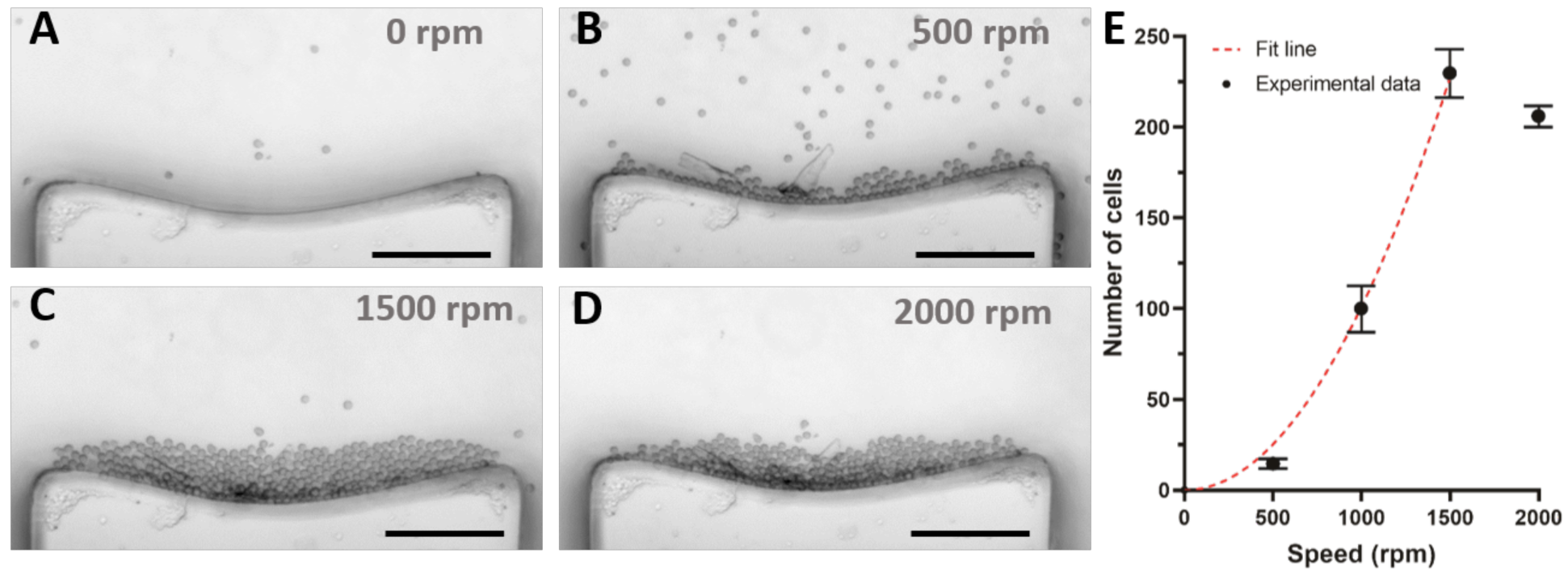
Cell alignment via centrifugation. (**A-D)** Effect of centrifugation speed on the alignment of the cells within the concave barrier. (Scale bar = 100 µm) (**E)** Number of aligned cells as a function of rotation speed (Error bars= mean ± standard deviation, *n* = 5 experimental repeats,*R*^2^ > 0.98).

### Chemotaxis Validation

To validate neutrophil chemotaxis in the microfluidic device, we performed chemotaxis assay in gradients of fMLP using human neutrophils. Neutrophils were isolated from whole blood using a commercial magnetic immunoselection kit and resuspended in a cell migration media. Cells were infused into the device after fibronectin coating and BSA blocking. After a short incubation period, a chemoattractant gradient was generated by pipetting fMLP solution into the source reservoir. Time lapse images of neutrophil migration were recorded on an inverted microscope and cell tracking performed through an ImageJ plugin (**Video S2**). Over 85% of cells in fMLP gradients had an elongated morphology in the first few minutes when compared to the rounded morphologies seen in the absence of fMLP gradient in control devices. Analyzing the cell tracks at an individual cell level revealed minimal migration in control devices (**Figure 4A**), but substantial directional bias of neutrophil migration in 100 nM fMLP gradients, with 80% of all cells moving towards the source of the chemoattractant (**Figure 4B**). In the 100 nM fMLP gradient, neutrophils had an average migration velocity of 2 µm/min and the velocity did not significantly change after 30 minutes (**Figure S2**). The cell paths were found to have an average directness factor of 0.3 (ratio of displacement to total distance), showing a strong preference for fMLP gradient. In contrast, the control sample showed no net displacement and did not possess a directional bias. We further assessed the variation of neutrophil migration speed as a function of time, which stabilized after ∼30 min as the gradient reached stable values (**Figure S2**). These results confirm that our microfluidic device is able to elicit a chemotactic response from the added cell sample.

**Figure 4.**
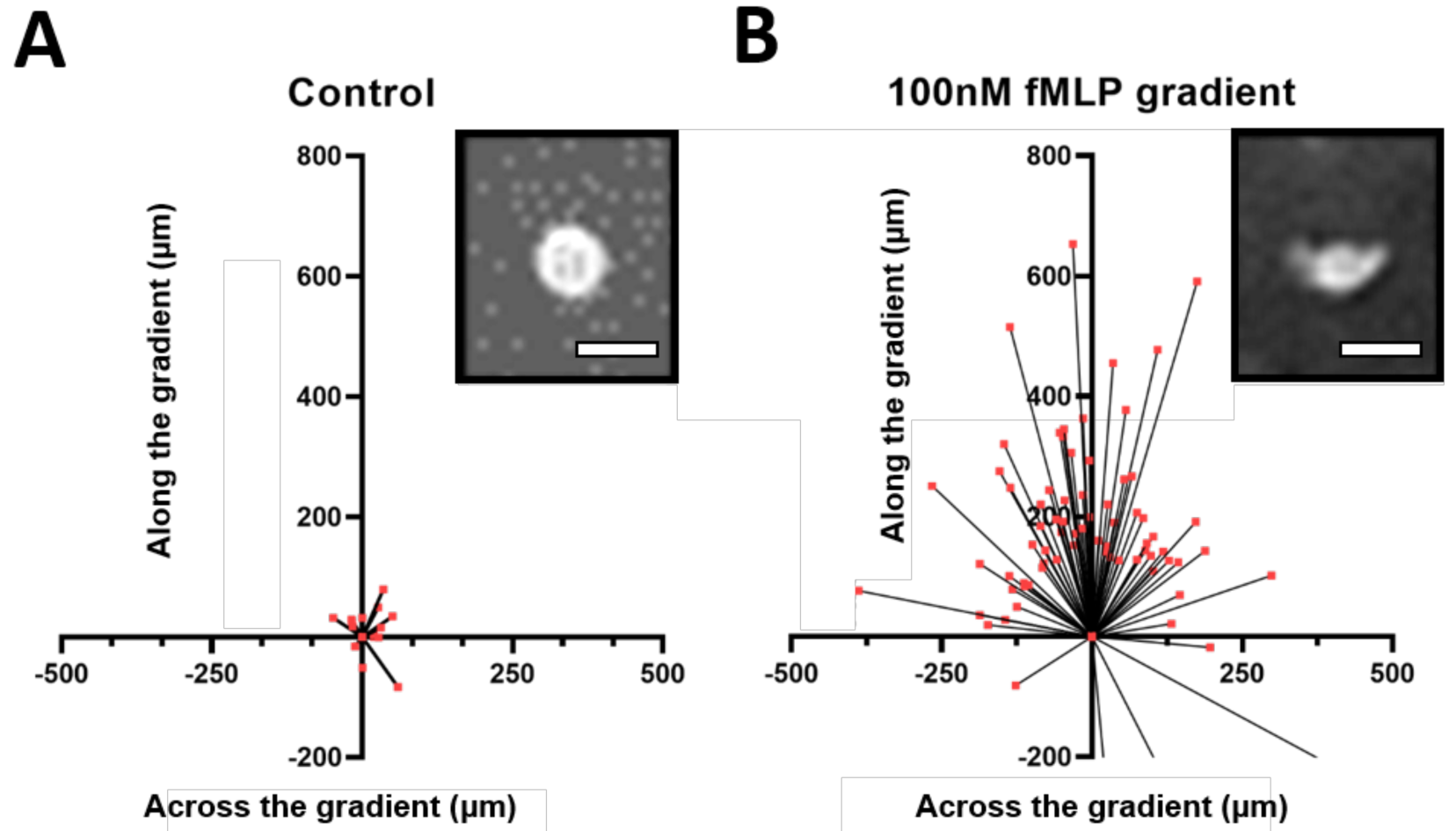
Validation of chemotaxis in the microfluidic device. (**A)** Real-time tracking of the initial and final cell positions after two hours of incubation in the absence of fMLP chemoattractant. **Inset:** Spherical cell morphology observed under these conditions. (**B)** Real-time tracking of the initial and final positions after two hours of migration in a 100 nM fMLP gradient. **Inset:** Elongated morphology adopted by actively migrating cells. (Scale bar = 10 µm)

### End-point Analysis Validation

To establish the effectiveness of the end-point chemotaxis assay, we repeated the neutrophil chemotaxis experiments with the additional step of centrifugal alignment. We determined the locations of cells, relative to the alignment barrier, by automatic segmentation of a microscope image of each channel after two hours of incubation at 37°C. The end-point chemotaxis assay was validated by comparing the final positions of neutrophils after incubation in devices with chemoattractant gradients created using various fMLP concentrations. In control devices without fMLP gradient, only a few cells were observed to migrate away from the barrier. In devices with fMLP gradient, most of the cells were observed to migrate away from the barrier (**Figure 5A-C**). Varying the fMLP gradient, we found that neutrophils appear to require a threshold fMLP gradient to activate chemotaxis since migration was observed only when the fMLP gradient was greater or equal to 25 nM (**Figure 5D-F**, p < 0.005). There also appears to be a decrease in migration distance from 25 nM to 100 nM, but this trend was not statistically significant (p > 0.05). Together, these data present support the use of centrifugal cell alignment as a rapid and convenient multiplexed end-point cell migration assay.

**Figure 5.**
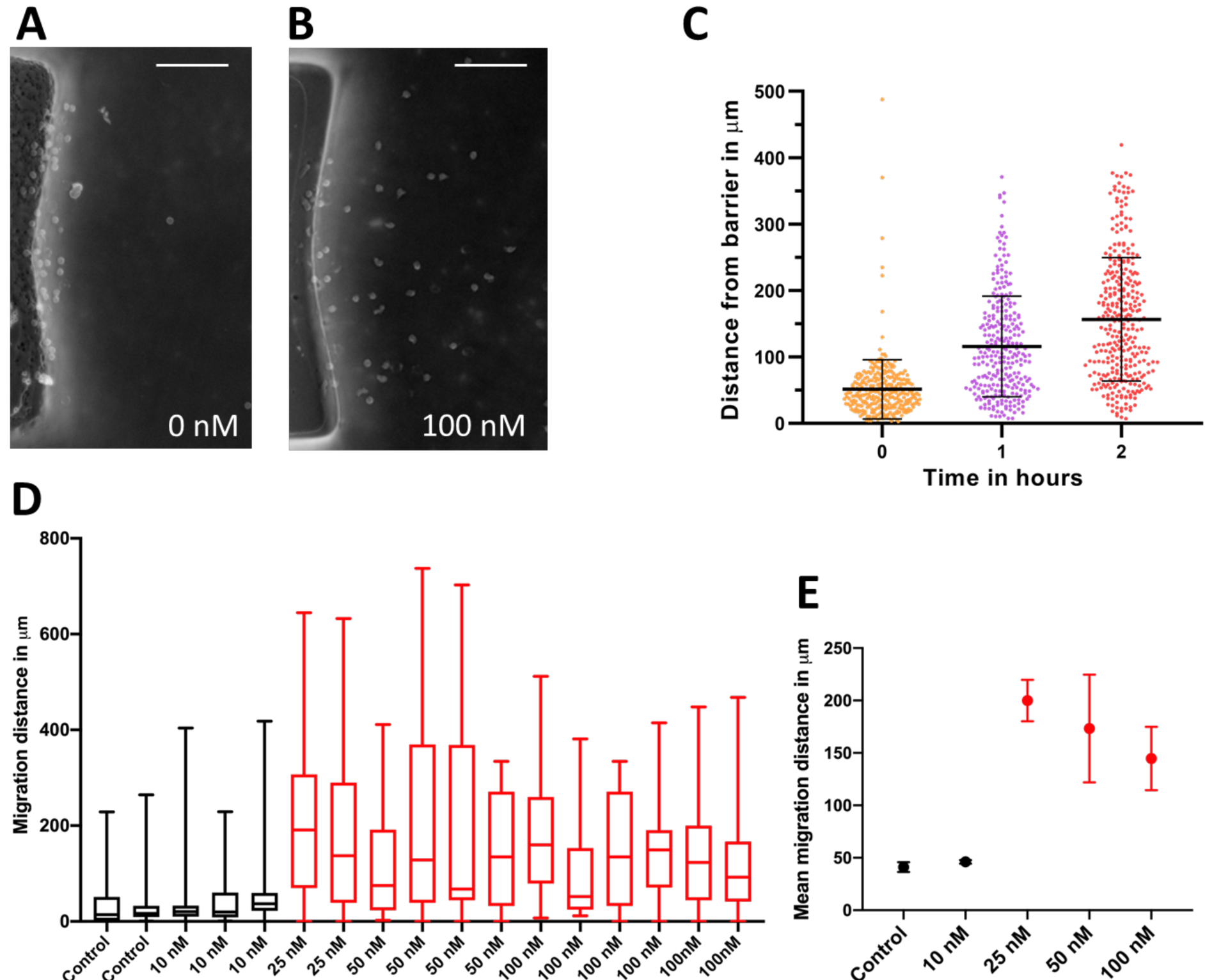
Endpoint analysis chemotaxis assay. (**A-B)** Micrographs of cells near the barrier features following alignment and two hours incubation for control (A) and a 100 nM fMLP gradient (B). (Scale bar = 100 µm) (**C)** Positions of cells along the length of the channel over two hours of migration in a 100 nM fMLP gradient (Error bars indicate standard deviation, n> 240). (**D-E)** Cell position distribution after chemotaxis as a function of fMLP gradient strength (Error bars indicate standard deviation). Observed differences in the mean of migration distance between ⩽10 and ⩾25 nM fMLP were statistically significant (p < 0.005).

## Discussion

In this study, we investigated an approach to develop a multiplexed end-point chemotaxis assay by aligning the cell sample against a barrier via centrifugation. The aligned cells are then exposed to a passively generated chemoattractant gradient, which causes the cells to migrate away from the barrier. Fluorescence imaging was used to observe chemical gradient profiles and validate its stability up to 3 hours while cell alignment at the barrier feature after centrifugation at various spin speeds was quantified through microscopy. Neutrophils in gradients of fMLP showed elongated morphologies and directed migration, while cells in control devices were rounded and non-motile. End-point analysis was used to obtain the average displacement of the cell population in response to the gradient as well as the spread in the final positions of cells after migration. After migration in an incubated environment, the chemotactic capability of the cell sample could be determined from a single microscopy image of the cells in the device.

Recent work on microfluidic chemotaxis assays often require real-time microscopy to track cell migration paths. While these experiments can be multiplexed using automated image acquisition, the requirement for cell tracking places fundamental limits on the throughput of these assays. Specifically, tracking fast moving cells, such as neutrophils, require acquiring an image every 30 seconds. The time required for moving the microscope stage and for image acquisition places a limit on the number of assays that can be analyzed in parallel. Experimental throughput can be increased by using advanced cell tracking algorithms that do not assign individual cell tracks, but instead derives aggregated populational measurements from the sample^21^. However, this approach does not fundamentally improve the scalability of these assays.

End-point chemotaxis assays that align the cell sample at a common starting point can dramatically improve the throughput and scalability of chemotaxis assays by providing a readout from a single microscopy image. Previous efforts to develop end-point chemotaxis assays used traps and sieves to position cells at a common starting point. The cells must be transported into these locations using carefully controlled fluid flow, which involves long preparation times and can require large volumes of reagents, and is therefore difficult to scale. Aligning cells using traps and sieves also applies significant shear stress on the cells, which can affect their migratory properties^26^. Additionally, the geometry of the traps or sieve must be matched to the geometry of the cell as well as the density of the cell sample, which makes this approach less robust when dealing with heterogeneous samples, such as primary samples from patients. Our centrifugal alignment method provides an alternative approach for cell alignment that does not require precise flow control, applies minimal shear stress to cells, and is largely insensitive to cell geometry. Cells can simply be pipetted into the device, where they are centrifuged to align against a concave barrier feature designed to minimize inertial shear. The motion of cells within the device is primarily driven by centrifugation speed, which can be easily adjusted based on the cell sample, device geometry, and surface adhesion forces. This approach is also compatible with heterogeneous cell samples where target cells can be fluorescently labeled and then identified by imaging after chemotaxis. Therefore, centrifugation-based alignment is an attractive option for developing scalable multiplexed end-point chemotaxis assays.

Finally, the microfluidic device design described here provides a number of favourable features that enable high-throughput experiments by scaling to hundreds of simultaneous chemotaxis assays in parallel. Specifically, the microfluidic device has a small footprint, which enables the integration of a large number of experiments on a single substrate. The design is a simple single-layer device that is easy to manufacture and does not require precise mechanical assembly. The device preparation and cell alignment does not require external controls for fluid handling thus leading to simple operation that can be automated using a pipetting robot. Each individual device requires only a small number of cells (500-5000 cells/ml) and reagents (5 µl). Centrifugation allows for simultaneous alignment of a wide range of cell types seeded at various densities. All these features result in a stand-alone multiplexed device that can be infused with reagents and cell samples and then simply be placed in the incubator for chemotactic migration. In conclusion, we demonstrate a simple and scalable multiplexed end-point chemotaxis where cells are aligned to a common starting point by centrifugation.

## Materials and Methods

### Preparation of Primary Neutrophils

Healthy donors between the ages of 18 and 70 were included in this study. Following informed consent and in accordance with University of British Columbia Research Ethics Board guidelines (UBC REB H10-01243), whole blood was collected into sodium EDTA tubes (BD Vacutainer). Neutrophils were enriched using a negative selection neutrophil isolation kit (Cat #19666, 18000, Stemcell Technologies, Canada) according to the manufacturer’s instructions. Neutrophils were washed and resuspended in RPMI-1640 (Cat #11875119, Gibco) with 0.4% Bovine Serum Albumin (BSA, Cat #37525, Gibco). The density of neutrophils was kept at ∼**10.4 × 10**^**6**^ /ml. To ensure evenly distribution of neutrophils in microchannels, the cells were pipetted up and down several times first and then were loaded from the sink reservoirs at a very low speed. All purified neutrophils were used within 2 hours after the separation.

### Device Fabrication

Our microfliudic device consists of 12 microchannels with two reservoirs (diameter 3 mm, height 2 mm) connected by a straight channel (length 2 mm, width 1 mm, height 0.3 mm). The alignment feature/barrier is 0.4 mm long, 0.3 mm wide and with a concavity on one edge. 3D printed molds of the device (Protolabs, Raleigh, USA) were used for device fabrication. The moulds were pre-treated at 65°C for 48 hours prior to use. PDMS silicone (Sylgard-184, Ellsworth Adhesives, Germantown, WI, USA) was mixed at a 10:1 ratio with the curing agent (Sylgard-184, Ellsworth Adhesives, Germantown, WI, USA) and poured into the molds. The molds were degassed and heat-cured for 2 hours at 65°C. The cured PDMS was removed from the molds and inlet holes for the source and sink reservoirs were punched using a 1 mm hole punch (Technical Innovations, Angleton, TX, USA). The device was then bonded to a 2” x 3” glass slide (TedPella Inc, USA) using air plasma (Cat #PDC-001, Harrick Plasma). The freshly bonded device was immediately filled with phosphate-buffered saline (Cat #10010023, Gibco) to prevent air bubble formation.

### Device Preparation and Cell Alignment

The plasma activated device was infused with 30 µl fibronectin (0.1 mg/ml) extracellular matrix (ECM) protein solution in PBS and incubated one hour before flushing with 200 µL of Phosphate Buffered Saline. Subsequently, the device was primed by infusion of 30 µl 0.4% BSA/RPMI, followed by incubation for one hour at room temperature. The device was then flushed with 30 µl fresh 0.4% BSA/RPMI, followed by seeding of 15 µl enriched neutrophils at the desired cell density. To align cells within the device, the inlets were then sealed with tape (Cat #96042, 3M) and the device was transferred to a Headway spinner at 2000 RPM for one minute, followed by incubation at 37°C for 15 minutes.

### Gradient Formation and Chemotaxis Assay

A chemoattractant solution was made by preparing 100 nM N-Formyl-Met-Leu-Phe (fMLP, Millipore Sigma) in 0.4% BSA/RPMI, supplemented, with 0.1% w/v FITC-dextran (Millipore Sigma). Immediately following cell alignment, 5 µL this fMLP solution was introduced into the source reservoir and a thin layer of silicone oil (Cat #378399, MilliporeSigma) was applied to cover the openings to minimize evaporation. For continuous cell tracking, the device was imaged with a Nikon Ti-2 microscope. The images were analyzed using the FIJI cell tracking plugin from ImageJ^29^. The cell paths obtained were later parsed and analyzed using custom Python scripts to visualize raw data. For end-point analysis, the microfluidic device was incubated in a 37°C humidified incubator for two hours. The device was subsequently imaged and cell migration distance was measured using ImageJ.

### Simulation of Gradient Formation

COMSOL Multiphysics was used to simulate and validate the gradient progression and evolution in the device. The simulation was established based on device geometry imported directly from the CAD file used for fabrication. The laminar flow profile and transport of diluted species physics modules were used to model the gradient of FITC Dextran (D = 6 ×^-6^*cm*^-6^*s*^-1^). No slip conditions were applied to obtain the initial profile of the gradient due to flow. The simulation assumed chemoattractant solution was introduced into the device at a flow rate of 1 µl/s for 5 seconds.

## Additional Information

### Authorship Contributions

H.M. conceived the idea and supervised the work. S.S., P.D., and K.M. performed the experimental work. All authors wrote the manuscript.

## Acknowledgments

This work was supported by grants from a NSERC (2015-06541, 508392-17) and CIHR (381129). P.D. acknowledges funding from the China Scholarship Council. K.M. acknowledges funding from MITACS Accelerate program (IT09621). H.M. acknowledges funding from the CIHR New Investigator Salary Award program (322375).

