## Supplementary material for "Multiplexed End-point Microfluidic Chemotaxis Assay using Centrifugal Alignment": SI

### Electronic Supplementary Information (ESI)

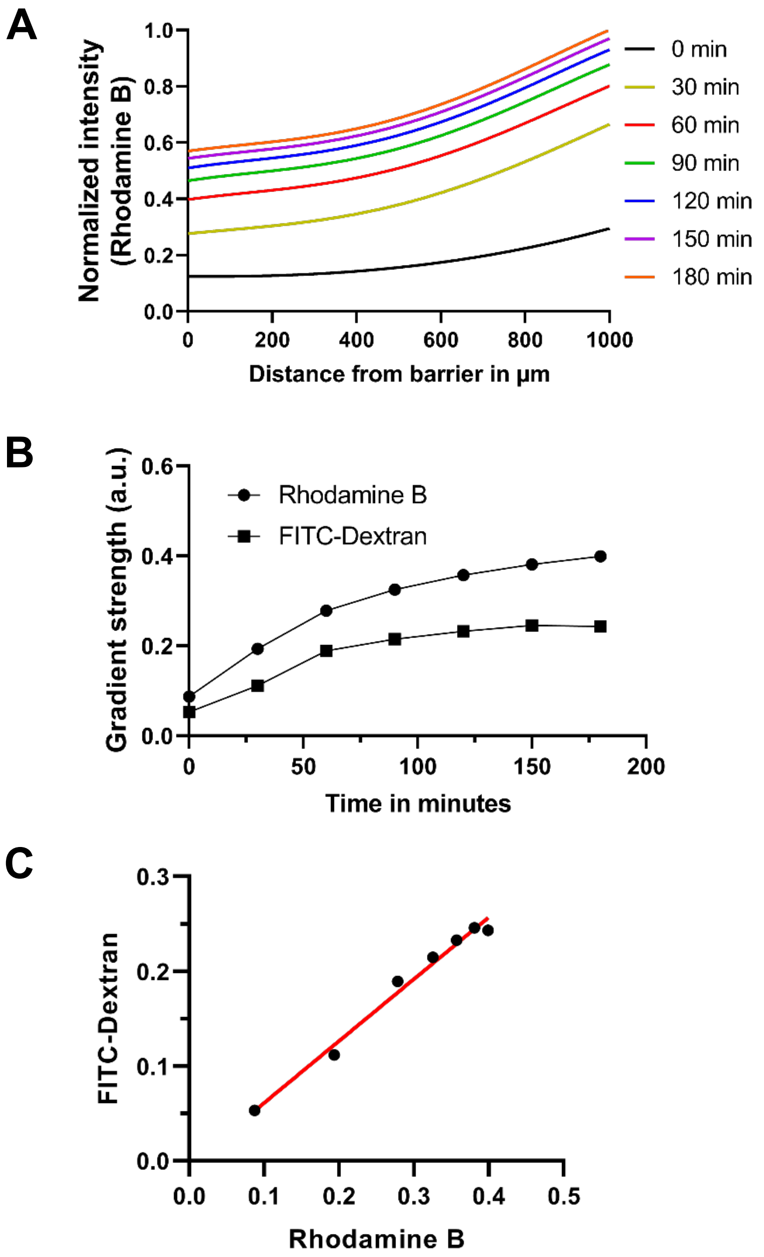


**Figure S1.** Experimental study of gradient formation in the design with the barrier feature. (A) The gradient profile was visualized using Rhodamine B, which has a similar molecular weight as fMLP. (B) The gradient strengths of Rhodamine B and FITC-Dextran measuring a function of time. (C) A regression line superimposed on the data from (B), showing a strong correlation between the gradient strength of Rhodamine B and fMLP-Dextran ($R^2 = 0.99$).

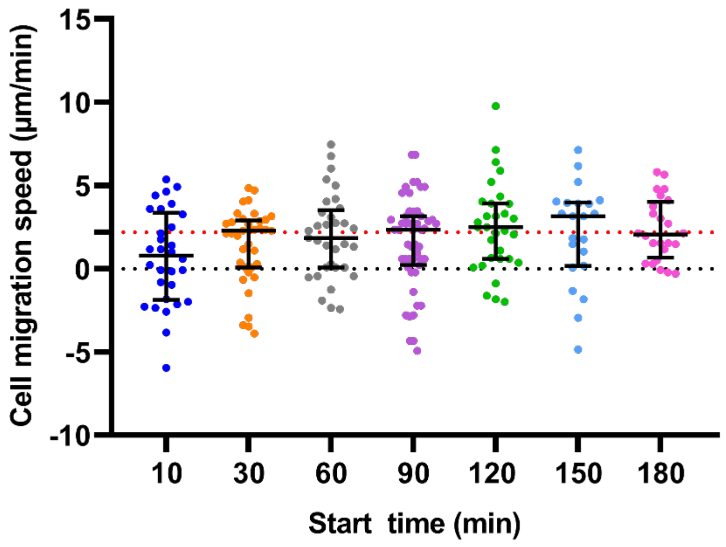


**Figure S2.** Average migration speed of neutrophils along the direction of microfluidic channels, measuring within 10 min after the start time. (N >30, error bars indicate median with interquartile range)

**Supplementary video 1:** Gradient formation using FITC-Dextran in the design with the barrier feature in 180 min.

**Supplementary video 2:** Human neutrophil migration in the presence of 100 nM fMLP gradient within 150 min.
